# Massive Phenotypic Measurements Reveal Complex Physiological Consequences of Differential Translation Efficacies

**DOI:** 10.1101/209098

**Authors:** Adam Paul Arkin, Guillaume Cambray

## Abstract

Control of protein biosynthesis is at the heart of resource allocation and cell adaptation to fluctuating environments. One gene’s translation often occurs at the expense of another’s, resulting in global energetic and fitness trade-offs during differential expression of various functions. Patterns of ribosome utilization—as controlled by initiation, elongation and release rates—are central to this balance. To disentangle their respective determinants and physiological impacts, we complemented measurements of protein production with highly parallelized quantifications of transcripts’ abundance and decay, ribosome loading and cellular growth rate for 244,000 precisely designed sequence variants of an otherwise standard reporter. We find highly constrained, non-monotonic relationships between measured phenotypes. We show that fitness defects derive either from protein overproduction, with efficient translation initiation and heavy ribosome flows; or from unproductive ribosome sequestration by highly structured, slowly initiated and overly stabilized transcripts. These observations demonstrate physiological impacts of key sequence features in natural and designed transcripts.

## INTRODUCTION

Timely production of functional proteins to express the most adequate phenotype in a given environment and at minimal biosynthetic cost drives the perpetuation of organisms and their underlying genetic information over time. Control of gene expression consists of two successive processes, transcription and translation, which are often naively treated as independent: transcriptional regulatory networks define the spatio-temporal coexpression of hard-coded DNA information into labile messenger RNAs molecules in a first step; while these informational intermediates enable controllable signal amplification as they are translated into proteins in a second step.

Much attention has been devoted to transcription control as a prime determinant of regulatory network dynamics. The significance of translation control of global regulation has only begun to emerge (Vogel and Marcotte, 2012). Translational processes play a decisive role in the cellular economy, with half the energetic expenditure of growing E. coli cells dedicated to protein biosynthesis and ca. 40% of that production devoted to making the translation machinery itself (Li et al., 2014). Consequently, cells have evolved remarkably simple and robust regulatory strategies to balance the production of ribosomes and other necessary proteins in response to varying growth conditions (Scott et al., 2014). These differential allocation patterns deeply constrain the proteome and the physiological state of the cell.

There have been a number of intriguing reports questioning the directionality of the central dogma. For example, a transcript’s abundance has been linked to its translation efficiency, through modulation of either transcription (Belogurov and Artsimovitch, 2015) or degradation rate (Hui et al., 2014). Precise characterization of such feedbacks is essential to refine our understanding of regulatory networks and improve functional predictions.

We implemented a library comprised of over 244,000 sequence variants of a fluorescent reporter gene designed to systematically explore combinations of the most prominent coding sequence properties believed to impact translation (Figure 1). This library is organized around 56 replicate series—each implementing a full-factorial design around tight and distinct foci in sequence space. Although varying only 96 nucleotides (nts) at the beginning of a gene controlled by otherwise constant regulatory signals, this design of experiment could at best account for a third of the resulting variance in protein production (companion paper).

**Figure 1.**
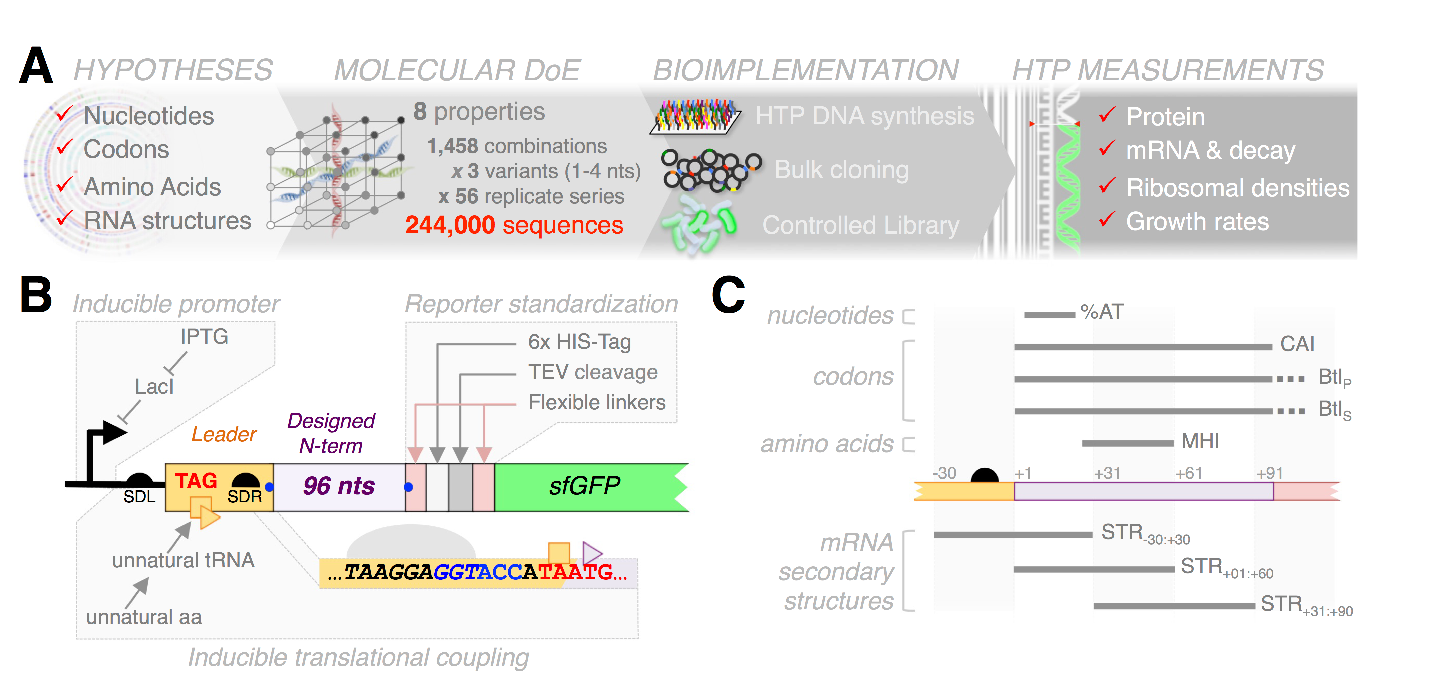
Molecular design of experiment at the genome scale. **(A)** High-throughput Design of Experiments at the molecular level. Sequences were designed to produce full-factorial libraries of 8 sequence properties correlating with protein production in E. coli. Designer sequences were synthesized and cloned in bulk. Various phenotypes are characterized in parallel using high-throughput sequencing as a generic quantitative readout. **(B)** A standard reporter for translational output. Synthetic sequences are cloned as N-terminal fusions to a modified sfGFP, which produces an invariant reporter upon post-translational processing by the TEV protease. Translation of the reporter is driven by a perfect Shine-Dalgarno motif (SDR, italics) embedded into a novel inducible translational coupling device. Secondary structures in the initiation region can be controllably disrupted by adjusting the influx of leader-bound ribosomes through tunable induction of an unnatural suppressor system. **(C)**Topology of the sequence properties varied in the factorial design.

To better understand the mechanisms underlying the variable observations, we complemented data on protein production with highly parallel measurements of cellular growth rate, reporter transcript abundance, decay and ribosome loading for virtually all strains in the library. The unique dataset resulting from these experiments permits unraveling of the mechanistic consequences of sequences variations on a core cellular process and their significance in impacting cellular fitness.

In this paper, we detail our design framework and dissect the impact of the designer sequence perturbations on protein production in different conditions of translational coupling. A more systemic analysis integrating these results with the other phenotypic responses is presented in a companion paper.

## Results

### Coding sequence perturbations reveal a biphasic relationship between protein production and cell growth

Gratuitous expression of heterologous genes imposes a global cost on the cell, which can result in severe growth defects. This burden reflects accrued consumption of nucleotides, amino acids and other metabolites, as well as the temporary mobilization of limited cell resources, such as tRNA and ribosomes (Ceroni et al., 2015; Scott et al., 2010; Stoebel et al., 2008). In theory, these complex physiological perturbations are strongly dependent on the interplay between translation initiation and elongation rates (Shah et al., 2013). Since the sequence variations engineered in our library affect both rates (companion paper), we suspected substantial physiological differences between strains. We used high-throughput sequencing of the designed sequences to derive parallel measurements of relative growth rates from bulk competition experiments within the entire library (Hietpas et al., 2011; van Opijnen et al., 2009).

We first propagated the library by serial dilution in the same rich, controlled medium as previously used for measuring protein production. For each strain, we estimated relative growth rate using the log-ratios of sequencing read frequencies measured in the propagated populations to those observed in the initial population. To ensure we had both the adequate read depth across the entire library while still obtaining a good dynamic range for growth rate estimation, we combined measurements of competitions after *ca.* 13, 28 and 60 generations (Material and methods; Figure S2ABCD). Based on three replicate competition assays at generation 60, we estimated a reasonably low experimental error for this type of batch competition assay (5% of the total variance). We were thus able to derive precise measurements of growth rate in regular conditions (W_NC_) for 233,846 strains.

As observed for protein production, the distribution of growth rates varies greatly between factorial series (Figure 2A). The sequence properties implemented in our molecular Design of Experiments (Figure 1C) and their second-order interactions account for only 5-31% of the variance in W_NC_, as quantified by ANOVA within each factorial series (14% on average; Figure 2B). The secondary structure across the start codon (STR_−30:+30_) and that designed farther in the coding sequence (STR_+31:+90_) show the most prevalent effects, accounting for 30% and 19% of explained variance on average, respectively. These structures have contrasting effects and sizeable interactions: weak STR_−30:+30_ negatively impact growth whereas weak STR_+31:+90_ has a positive effect, especially when associated with strong STR_−30:+30_ and weak STR_+01:+60_ (Figure S2E).

**Figure 2.**
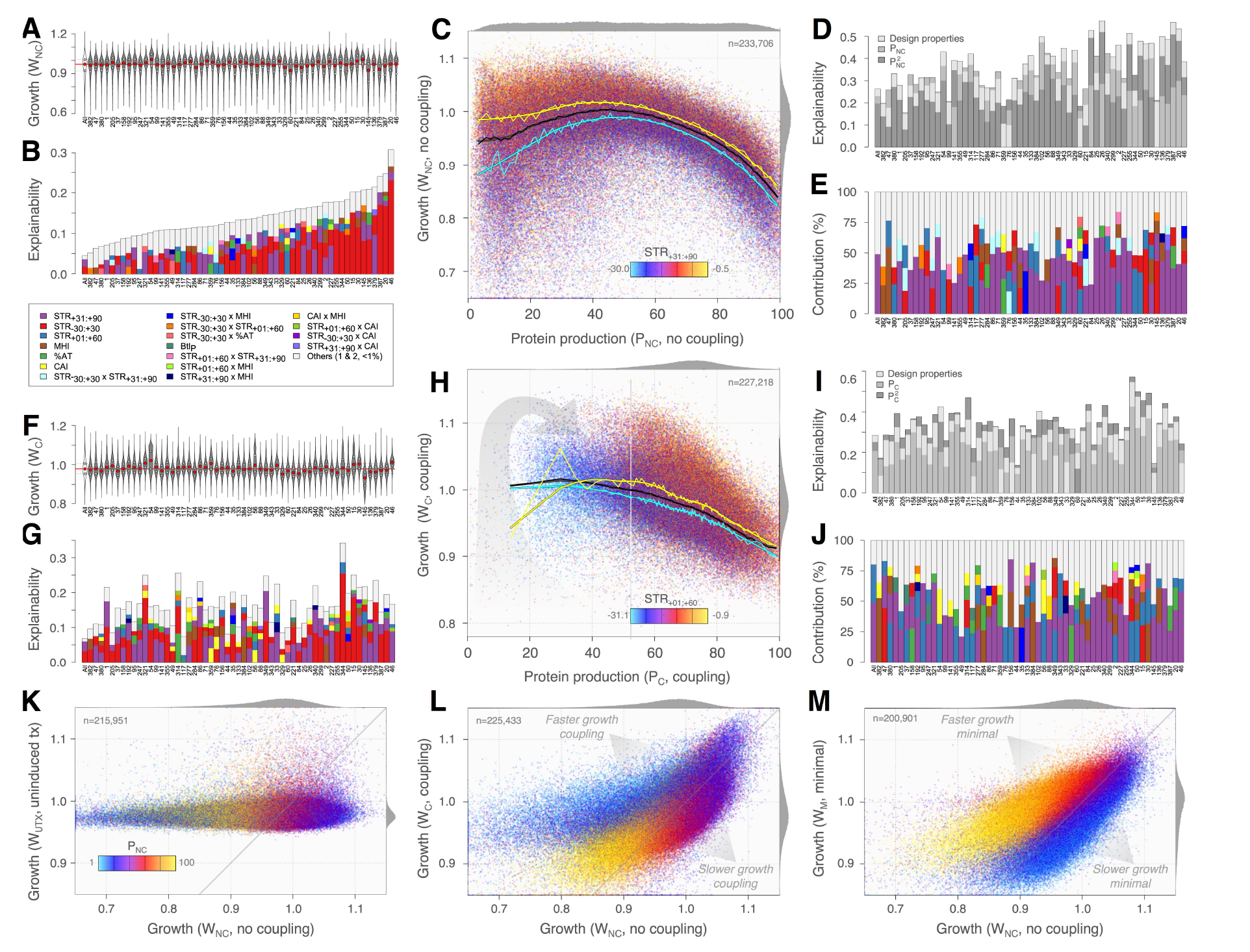
Unexpected growth defects associated with low translation initiation. **(AB)** Highly variable growth profiles and corresponding patterns of explanation between replicate factorial series. Violin plots in **(A)** show the distribution of growth rate under non-coupling condition (W_NC_) for each series. Red and grey dots mark medians and interquartile ranges, respectively. Stacked bars in **(B)** show corresponding variance decomposition by ANOVAs based on the sequence design properties. Effects≥1% are sorted in decreasing order from bottom to top and colored as shown. Other effects are grouped on top in grey. **(CDE)** Unexpected relationships between growth, protein production and within-genes mRNA structures. Scatter plot of W_NC_ versus P_NC_ colored by STR_+31:+90_, as shown **(C)**. The dark lines mark the median W_NC_ for each percentile of P_NC_ (thin line) and show a loess smoother highlighting the trend (thick line). Low P_NC_ is associated with W_C_, resulting in a strongly biphasic relationship. Yellow and cyan lines show the top and bottom deciles of STR_+31+90_, respectively. STR_+31+90_ is positively correlated with W_NC_, especially at low P_NC_. A multiple linear regression of W_NC_ including P_NC_ and P_NC_^2^ as explanatory variables improves explanatory patterns **(D)**. The importance of the squared component underscores non-linearity in most series. Bars in **(E)** detail the relative contributions of design properties in this regression and confirm the prominent impact of STR_+31:+90_ on W_NC_ across series. **(FGHIJ)** Same analyses under translational coupling. Epigenetic improvement of initiation causes substantial linearization of the relationship between PC and W_C_ **(HI)**. The impact of STR_+31:+90_ is largely maintained **(J)**. **(KLM)** Low rates of translation initiation are linked to slow growth. Scatter plots of growth rates in different culture conditions against the reference W_NC_, colored by P_NC_ as shown. Expression of reporter transcripts is necessary to observe growth differences **(K)**. Improving initiation by translational coupling increases growth rates amongst lowest protein producers **(L)**. Lower ribosome availability upon growth in minimal media increases the growth defect in low protein producers **(M)**.

Perhaps explaining the modest performance of this linear model, our data show a strongly non-monotonic relationship between W_NC_ and protein production (P_NC_, Figure 2C). The expected anti-correlation is confined to higher production, whereas lower producers demonstrateunexpected spread toward low growth. This relationship is generally well captured by quadratic regressions of P_NC_ on W_NC_ (1–52% of the total variance across series; 28% on average, with 19% by the squared component only; Figure 2D).

These regressions expose direct contributions to growth that are not mediated by effects on protein production. On average, design properties and their interactions explain 10% of the variance in W_NC_ not accounted by P_NC_ (Figure 2D). The contribution of STR_−30:+30_ to W_NC_ is then reduced to only 11% of the explained residual variance (Figure 2E). The apparent effect of this structure on W_NC_ is therefore largely due to its proximal impact on translation initiation, whereby weaker structures enable higher protein production and consequent fitness costs. In contrast, the respective contributions of STR_+31:+x90_ and STR_+01:+60_ respectively rise to 29% and 10% in this context, with weaker structures associated with higher growth in both cases (Figure 2E). This suggests a role for these structures in impacting cellular fitness that is distinct from their effect on P_NC_ (Figure 2C and (Figure S2GH). The larger spread of growth distributions amongst low protein producers further suggests strengthened contributions under initiation-limiting conditions (Figure S2EGH).

### Manipulation of expression uncover distinct costs associated to translation

We performed additional competitions in different growth conditions to clarify how these associations relate to the expression of the reporter (Material and methods). We first propagated three control populations for ca. 60 generations without inducing transcription of the reporter. As expected, the growth rates observed in this condition (W_UTX_) are much less variable and represent only 5% of the variance quantified above (Figure 2K). These results establish the necessity of transcription to the selection pattern observed previously.

Our reporter system is equipped with an inducible translation-coupling device capable of mitigating the adverse effect of mRNA secondary structure on initiation (companion paper and Figure 1B). To determine the effect of increasing initiation rates, we conducted competition experiments for populations propagated for ca. 26 and 33 generations under coupling and could derive W_C_ for 229,224 strains.

Improving initiation rates through elimination of mRNA structure around the Shine-Dalgarno motif decreases the variance in growth rate by about two thirds, while generally maintaining the relative growth profiles differences between individual series (Figure 2AF). ANOVAs on W_C_ globally provide a comparable picture to that obtained for W_NC_, with dominant effects of STR_−30:+30_ and STR_+31:+90_ (28% and 26% of the explained variance on average, respectively; Figure 2G). W_C_ and protein production (P_C_) display a drastically more linear relationship under coupling, demonstrating that the phenomenon of low initiation is in itself a key determinant of the fitness defects observed at low P_NC_ (Figure 2HL). Only 0–15% variance in W_C_ is accounted for by the squared component in quadratic regressions involving P_C_ (Figure 2I). Design properties then explain 3-24% of the residual variance, again with a large dominance of STR_+31:+90_ and STR_+01:+60_ (28 and 14% of explained variance on average, respectively; Figure 2J). Consistently, the few strains that are refractory to coupling due to overly strong STR_+01:+60_ (companion paper) remain associated with an inflexion toward lower growth (Figure 2H). Hence, low initiation rates as determined by structures in the translation initiation region (TIR) are necessary to the mechanism causing unexpected growth defects at low protein production.

To further probe the link between growth and translational regime, we sought to reduce the translation capacity of the cells. Bacteria grown in media with less nutrients are known to grow slower and produce fewer ribosomes (Schaechter et al., 1958). We thus propagated our library in minimal media for ca. 33 generations and could derive W_M_ for 201,893 strains. Low protein producing strains systematically exhibit lower WM for a given W_NC_ (partial correlation ρ=0.68, Figure 2M). The initiation-limiting properties of strong STR_−30:+30_ are no longer associated with a positive impact on growth (Figure S2F). The pattern of interaction with other structures is nonetheless maintained, such that strong STR_-30:+30_ combined with strong STR_+31:+90_ but weak STR_+01:+60_ determine lowest W_M_ (Figure S2F). These observations expose a component of the cost associated with translation that: i) is mechanistically distinct from the actual level of protein production; ii) is dependent on strong transcript folding; and iii) is amplified when fewer ribosomes are available.

### Differences in mRNA abundance are largely shaped by differential decay

In our system, transcription is driven by a constant promoter whose start site is located *ca.* 100 nts upstream of the designed sequences. Although we cannot rule out occasional interactions between designed sequences and transcription machinery, we do not expect extensive variations in transcription rates across strains (Kosuri et al., 2013; Mutalik et al., 2013). Instead, systematic differences in steady-state mRNA levels could arise from differential degradation kinetics arising from the sequence variations. Multiple lines of evidence indicate that translation rate and transcript degradation are intricately linked (Deana and Belasco, 2005; Huch and Nissan, 2014). Accordingly, coding sequence properties such as codon usage bias and mRNA folding have been associated with mRNA decay (Boël et al., 2016; Hui et al., 2014; Lenz et al., 2011; Presnyak et al., 2015).

We developed a targeted sequencing assay to follow the relative abundances of reporter transcripts after transcription was arrested *in vivo* (Figure 3A). Using this procedure, we could estimate steady-state RNA abundance (RNA_SS_), RNA half-lives (RNA_HL_) and an RNA protection index (RNA_PTX_) for 233,487 constructs (Material and Methods and Figure S3ABC).

**Figure 3.**
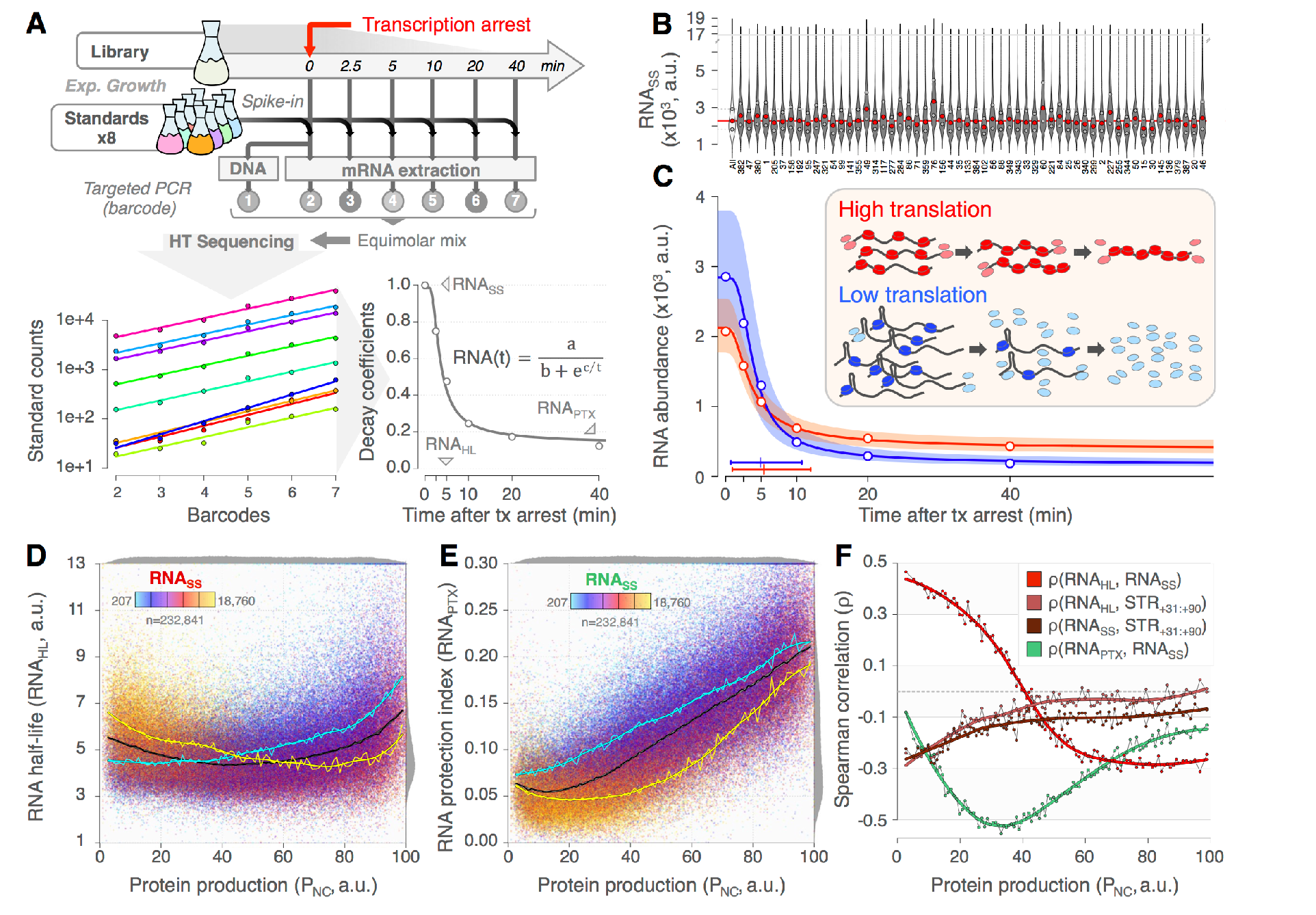
Translation rates affect RNA abundance. **(A)** High-throughput assay of RNA decay. Constant quantities of standard strains are spiked-in library samples. Standard sequencing reads increase as reporter transcript decay, which defines corrective coefficients. Corrected time series for each strain are fit to an exponential decay model to estimate RNA abundance at steady state (RNA_SS_; t=0), transcript half-life (RNA_HL_) and the final fraction of protected RNA (RNA_PROTECT_; t=+∞). **(B)** Diversity of RNA_SS_ profiles between replicate series. Red and grey dots mark medians and interquartile ranges, respectively. **(C)** Protein production impacts RNA abundance and decay. Red and blue points show medians for the top and bottom deciles of P_NC_, respectively. Red and blue lines and shadings mark corresponding decay fits and interquartile ranges. Lines on the bottom left show medians and interquartiles range of RNA_HL_. High protein producers are characterized by lower RNASS but higher RNAPTX. Ribosomal protection intensifies during the assay in highly translated transcript (inset, red). Secondary structures limit ribosomal protection but increase transcripts stability and RNASS (inset, blue). **(DEF)** Decay assay artifacts expose complex interactions between degradation and translation. Scatter plot of RNA_HL_ **(D)** and RNA_PTX_ **(E)** versus P_NC_, colored by RNA_SS_ as shown. Dark lines mark median RNAHL or RNAPTX for each percentile of PNC and the corresponding loess smoothers. Yellow and cyan lines show the top and bottom deciles of RNASS, respectively. **(F)** Various correlations for each percentile of PNC, as shown. Positive association between RNAHL and RNASS is restricted to low PNC and progressively inverted, reflecting swifter protection of fast initiated transcript (see panel C, red inset). RNAPTX is linearly related to PNC and modulated by RNASS. Stronger structures are associated with increased RNAHL and RNASS, especially at low PNC.

This experiment revealed substantial variability in RNA_SS_ over the whole library (CoV=0.44), as well as marked differences between factorial series (Figure 3B). Estimates of RNA_HL_ are equally variable though less reproducible (CoV=0.41, (Figure 3C and (Figure S3C), with a median half-life of 4.8 min typical of *E. coli.* Although longer-lived transcripts should be more abundant, the association between RNA_HL_ and RNA_SS_ is overall very poor (r=0.06, (Figure S3D). Closer examination reveals that decay profiles are strongly dependent on protein production Figure 3C). We identified a non-monotonic relationship between RNA_HL_ and P_NC_, such that transcripts with highest apparent stabilities are partitioned to the tails of the protein distribution Figure 3D). In this context, the correlation between RNA_HL_ an1d RNA_SS_ decreases smoothly with P_NC_ from *ca.* 0.4 to *ca.* -0.3, restricting the expected positive association to the lower third of protein producers (Figure 3F and (Figure S3D). These results suggest that distinct mechanisms are determining decay rates under extreme translational regimes.

The fraction of RNA_SS_ asymptotically left after decay (RNA_PTX_) is also strongly correlated with P_NC_ (r=0.76, Figure 3E). We initially introduced this parameter because it afforded better decay fits, but its biological significance was unclear. Our observation implies that individual transcripts are protected from complete degradation to an extent that depends on their effective translation rate (Figure 3C, inset). We have shown that the competition between ribosome binding and TIR folding is the chief determinant of initiation rates (companion paper). After transcription is stopped, the quantity of labile mRNAs decreases quickly, while that of stable t- and rRNA should remain approximately constant on the timescale of the assay. Thus, the relative titer of free ribosomes increases during the decay assay, promoting gradually improved initiation of the remaining transcripts (Bulmer, 1991; Shah et al., 2013). In this dynamic process, weaker structures are outcompeted faster (r=0.37 between RNA_PTX_ and STR _30:_+_30_ structure), resulting in larger protection amongst high protein producers.

Strains showing higher RNA_SS_ undergo slower and eventually lower protection (r=-0.39,Figure 3C and Figure 3E). Distributing a finite quantity of ribosomes over more transcripts leads to lower translation of individual transcripts (see Figure 5D), hence to less protection for longer times during the decay assay. Thereby, the apparent half-lives estimated for high protein producers capture the swiftness of the protection process during the assay rather than the stability of the transcripts *per se*— hence the negative correlation between RNA_SS_ and RNA_HL_ at high translation regime (Figure 3DF and Figure S3D).

**Figure 4.**
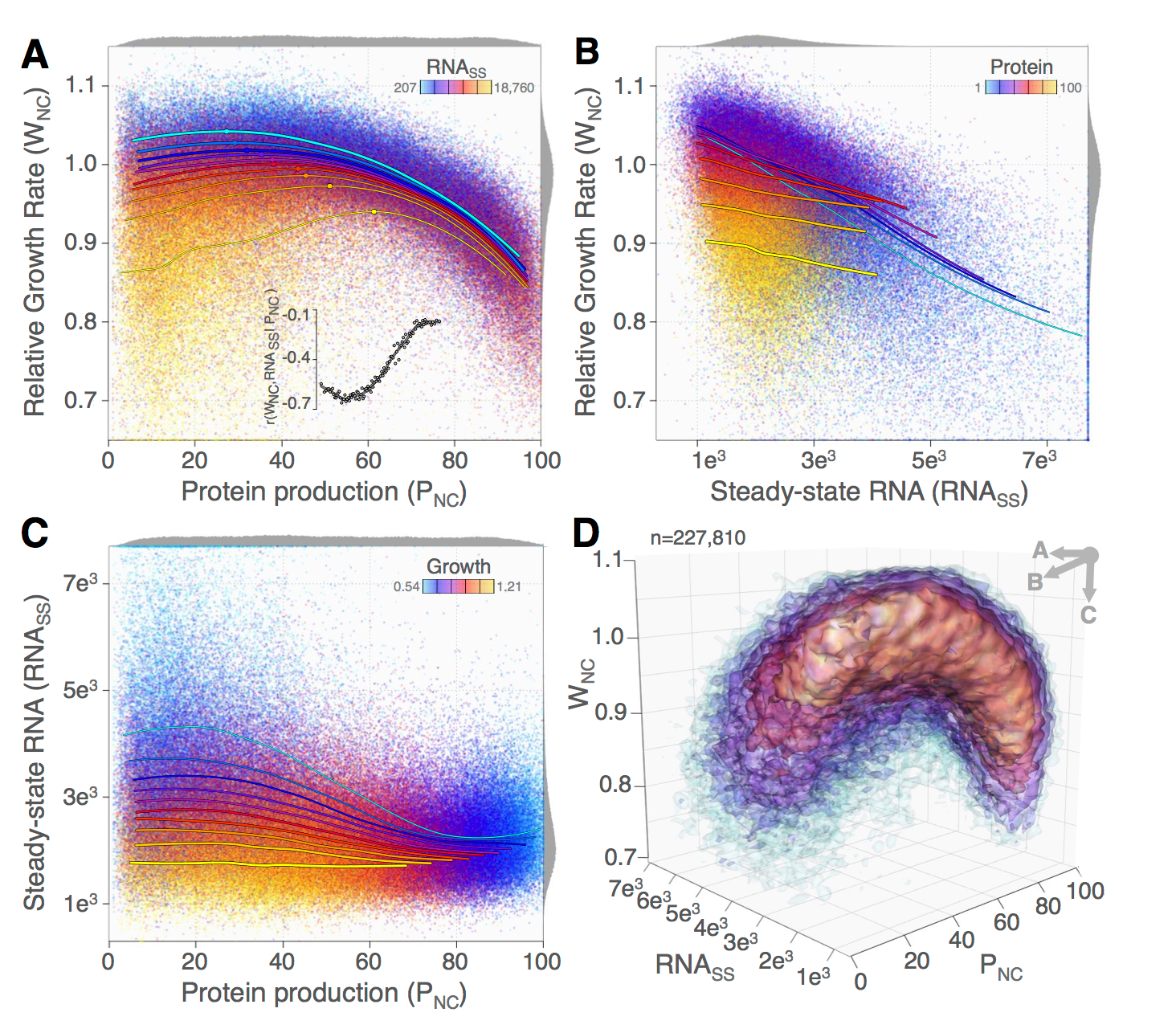
Slow growth caused by pathological cu-mu-lation of stable transcripts at low translation regime. **(ABC)** Scatter plots of two measured phenotypes, colored by a third as shown. Lines show loess regressions for every decile of the latter quantity, following the same color code. Line widths provide perspective corresponding to viewpoints in **D**. The biphasic relationship between W_NC_ and PNC in **A** mirrors that between RNA_SS_ and P_NC_ in **C**. It stems from the strong negative correlation between W_NC_ and RNA_SS_, which decreases with PNC (**B** and see inset in panel **A** for every percentile of P_NC_). **(D)** Three-dimensional phenotypic envelope of the library. Colored layers mark increasing data densities. Top-right arrows show viewpoints for panels **ABC**.

**Figure 5.**
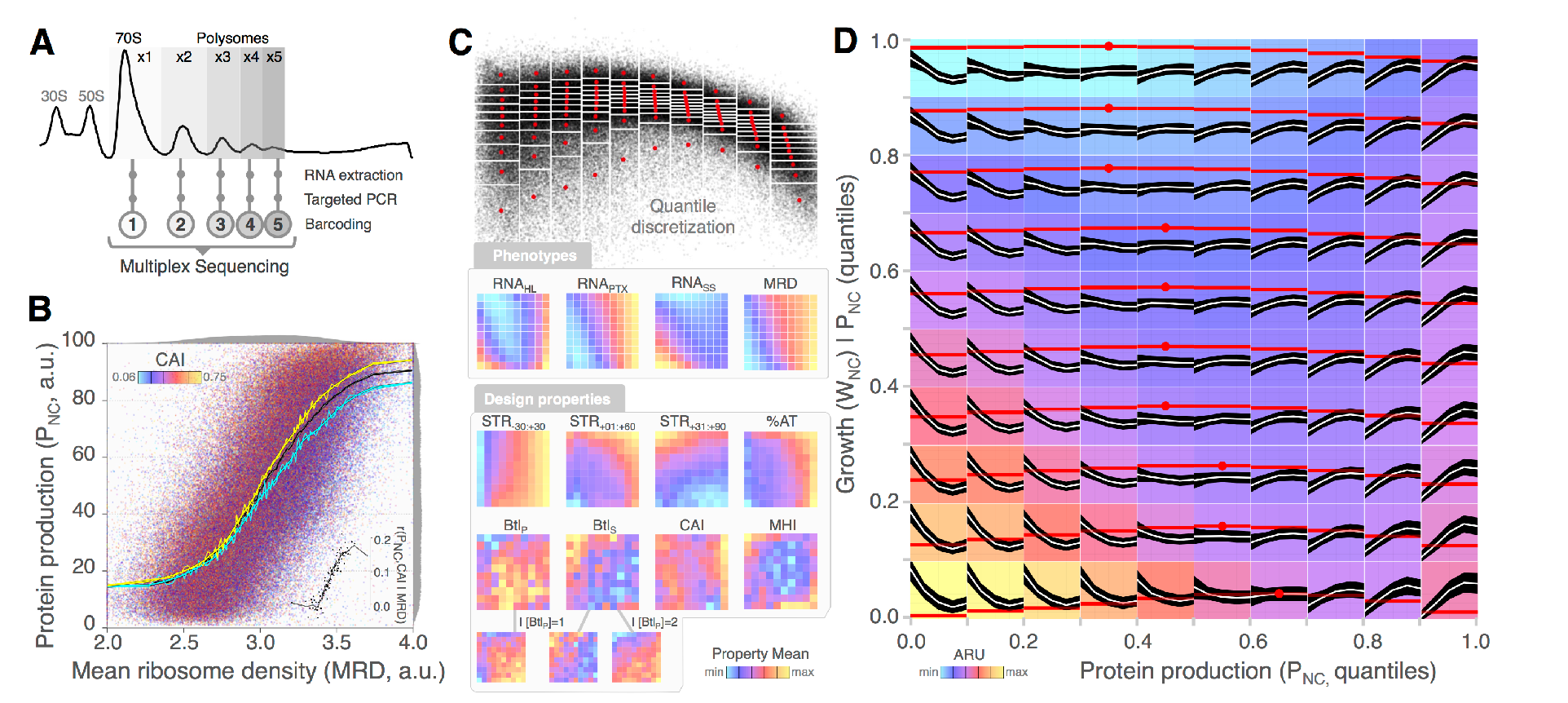
Ribosomal density profiles identify tipping points of translation cost. **(A)** High-throughput targeted polysome profiling. The first five polysome fractions were extracted from a sucrose gradient, barcoded and sequenced in multiplex. **(B)** Codon usage modulates protein production fro a given ribosomal density. Scatter plot of P_NC_ versus the mean ribosomal density (MRD), colored by CAI as shown. The dark lines mark the median P_NC_ for each density percentile (thin line) and a loess smoother highlighting the trend (thick line). Yellow and cyan lines show the top and bottom deciles of CAI, respectively. The correlation between CAI and P_NC_ increases with very percentile of ribosomal density (inset). **(C)** Coarse-grained grids of W_NC_ versus P_NC_. *Top:* Data are binned first by deciles of P_NC_ and then by deciles of W_NC_. Each bin comprise ~2,250 strains. Red crosses mark local averages. Bottom: Bins are color-coded to show the range of means in quantity indicated in each vignette. **(D)** Polysome profiles vary smoothly in this phenotypic space. White lines and dark shadings show median and interquartile ranges of the relative abundances of polysome fractions, from monosome (left) to pentasome (right). Red lines and points mark mean W_NC_ and corresponding row-wise optima. Background is color-coded by the apparent ribosome utilization (ARU, MRD times RNA_SS_). Increased prevalence of monosomes is associated with lower W_NC_ for a given P_NC_ (column, top to bottom). Row-wise W_NC_ optima correspond to balanced initiation versus elongation (flat profile).

### Dual dependence of RNA decay on translation regime and secondary structures

Transcript abundances are seemingly shaped by the antagonistic actions of two distinct translation-dependent mechanisms on decay, whose complex interplay eventually determines a biphasic relationship between P_NC_ and RNA_SS_ (Figure 4C).

At low initiation, transcript stability is largely determined by secondary structures that can hinder endonucleolytic cleavages by RNase E (McDowall et al., 1995) and processive degradation by 3’ exoribonucleases (Hui et al., 2014). Because both STR_−30:+30_ and STR_+01:+60_ are primary determinants of initiation rates (companion paper), their impacts on stability are confounded and challenging to discriminate. The effect of STR_+31:+90_ is more apparent, especially in the low translation regime (Figure 3F and S3EF)

Ribosomes are intrinsically capable of unfolding transcripts during initiation (Duval et al., 2013) and subsequent elongation (Qu et al., 2011), with rates commensurate to the thermodynamic stabilities of the structures. These processes likely promote degradation through transient exposure of single-stranded RNA substrates to ribonucleases. Accordingly, the dependence of RNA_HL_ and RNA_SS_ on secondary structures diminishes with increasing P_NC_ (Figure 3F), resulting in lower RNA_SS_ at intermediate translation regimes (Figure 4C) (Duval et al., 2013) and subsequent elongation (Qu et al., 2011), with rates commensurate to the thermodynamic stabilities of the structures. These processes likely promote degradation through transient exposure of single-stranded RNA substrates to ribonucleases. Accordingly, the dependence of RNA_HL_ and RNA_SS_ on secondary structures diminishes with increasing P_NC_ (Figure 3F), resulting in lower RNA_SS_ at intermediate translation regimes (Figure 4C).

At higher translation rates, heavy ribosome flows can compete kinetically with ribonucleolytic attacks (Boël et al., 2016) and particularly with the decay-initiating action of RNase E (Braun et al., 1998). Although blatant in the decay data (Figure 3E), this ribosome protection only results in a modest increase of RNA_SS_ amongst the highest protein producers during exponential growth (Figure 4C). This discrepancy probably reflects the limiting nature of ribosomes in growing cells, which is progressively relieved during the decay assay (Figure 3C, inset). The negative correlation between RNA_SS_ and RNA_PTX_ strongly supports this view (Figure 3EF).

The dual impact of folding on stability and translation gives rise to intricate functional interactions between designed structures (Figure S3G). For example, the stabilizing effect of strong STR_+31:+90_ is particularly marked when combined with strong STR_−30:+30_ (which effects low initiation) and weak STR_+01:+60_ (which minimizes thermodynamic competition with the two other structure). Similarly, the association between strong STR_+01:+60_ and lower transcript protection is more profound when initiation is not otherwise limited by potent STR_−30:+30_ and its folding not competed by strong STR^+31:+90^.

### Variations in RNA abundance impact cell growth

Variations in RNA_SS_ are strongly anti-correlated with W_NC_ (r=−0.45, Figure 4AB). Reflecting the dependence of RNA_SS_ on translation (Figure 4C), the strength of this association declines smoothly with P_NC_ (Figure 4A, inset). The co-distribution of these three phenotypic measures outlines a well-defined envelope that captures their constrained interdependences (Figure 4D and S4). The highest RNA_SS_ are found amongst the lowest P_NC_ and appear to determine the unexpectedly low W_NC_ observed in those strains. The variability in RNA_SS_ narrows down with increasing P_NC_ and its apparent contribution to growth defects decreases. Instead, W_NC_ becomes increasingly dominated by the cost of protein production. The cost associated with RNA_SS_ appears to be amplified at lower P_NC_, as initially noted for the causative STR_+31:+90_ (Figure 2C).

The transition from low to medium P_NC_ is generally accompanied by an increase in W_NC_, despite the added cost of accrued protein biosynthesis. Sequence variants with very low translation rates exhibit strong STR_−30:+30_ and/or STR_+01:+60_ *in vivo.* Since these features are also associated with increased transcript stabilities in these absence of translating ribosomes, the occurrence of sequence determining low RNA_SS_ is inherently sparser amongst lower than more intermediate protein producers. Given the apparent cost of RNA_SS_, this mechanistic contingency reduces the median growth rate observed at low protein production regimes. Nevertheless, fitness optima still lie at intermediate translation regimes when variations in RNA_SS_ are controlled for see Figure 4A, points on regression lines). The coordinates of these optima are tightly linked to RNA_SS_, with higher RNA_SS_ associated with lower optimal W_NC_ at higher P_NC_.

### Initiation-based ribosome sequestration help disentangling cellular labor from material costs

The observations above suggest the existence of a negative interaction between the physiological cost of high mRNA abundance and that of slow initiation. Since the sequestration of nucleotides and phosphate in polymerized transcripts is benign to *E. coli* cells grown in rich medium (Stoebel et al., 2008), high RNA_SS_ is unlikely to be the proximal cause of the observed costs. However, high heterologous transcript abundance can have dramatic consequences on the allocation of limited ribosomal resources toward the host transcriptome (Shah et al., 2013), eventually leading to decreased growth rates (Scott et al., 2010). Such differential ribosome mobilization by variant library transcripts provides an attractive mechanistic basis for the observed associations.

To examine this, we performed a deep-sequencing-based polysome profiling to quantify the distribution of ribosomes densities associated with each transcript variant (Figure 5A). We derived the relative enrichment of individual constructs in each of the first five polysome fractions and obtained a polysome profile for 240,403 strains (Material and Methods).

Although the shape of such profiles conveys important information regarding the translation kinetics of a given transcript, it is difficult to analyze at scale. We first conflated the data into a single measure that captures the average number of ribosome per reporter transcript. This mean ribosomal density (MRD) is highly correlated with P_NC_ (r=0.73), so that every ribosome density corresponds to a tight range of protein production (Figure 5B). Two sequence properties—CAI and STR_−30:+30_—show interesting signals in this context.

For a given density, P_NC_ increases with CAI (Figure 5B). The strength of this association smoothly increases with MRD, as translation becomes less limited by initiation (inset). This result is consistent with the expected effect of rarely used codons in slowing down elongation rates as a consequence of low abundance of cognate tRNA species (Bulmer, 1991). Increasing ribosome densities further reveals a strengthening association between CAI and RNA_PTX_ (up to r=0.17, Figure S5A) that underpins a similar though weaker correlation with RNA_SS_ (up to r=0.11, Figure 5SB). These data support the involvement of heavy and smooth ribosome trafficking in transcript protection.

Stronger STR_+31:+90_ is associated with increased P_NC_ for a given MRD, especially at medium densities (up to r=-0.22, Figure S5C), while it should instead slow elongation down and determine lower P_NC_, as observed for low CAI. To resolve this discrepancy, we consider the converse relationship, whereby strong STR_+31:+90_ is associated with increasingly lower densities as P_NC_ diminishes. This pattern almost perfectly mirrors the association of STR_+31:+90_ with RNA_SS_ (Figure S5D). Now, higher RNA_SS_ is globally associated with lower loads on individual transcripts (r=−0.31). It is therefore likely that the apparent effect of STR_+31:+90_ on MRD is an indirect consequence of the intricate, translation-dependent effects of secondary structures on RNA stability and abundance (Figure S5E). These observations illustrate the difficulties in untangling causality amongst many dependencies existing in the data.

To better characterize polysome profiles, we binned the library into equally sized subpopulations covering the extent of the protein-growth phenotypic space (Figure 5C). We then derived a statistical representation of the ~2,270 profiles in each bin (Figure 5D). At low P_NC_, profiles are dominated by low fractions characteristic of low initiation. As P_NC_ increases, profiles become increasingly flatter and progressively shift toward larger representation of dense fractions indicative of limiting elongation. Strikingly, the flat profile that marks the equilibrium between initiation and elongation always corresponds to local fitness optima (Figure 5D, red dots). These results further confirm the physiological cost associated with slow initiation.

The cost of initiation is most striking amongst strains with low P_NC_ and high RNA_SS_, whose profiles show highest skew toward low polysome fractions. Sequence variants confined to this phenotypic region are characterized by strong structures encompassing the TIR and downstream coding sequence (Figure 5C). Individually, these properties provide the requirements for slow initiation and improved transcript stability. We propose that their combined effects define a self-reinforcing catastrophic cycle whereby initially minor transcript stabilization spirals into a generalized translation decrease through a pathological transcript buildup driven by feedbacks between translation and stability (see Discussion and Figure 6F). The resulting convolution of high RNA_SS_ with slow initiation lead to the unproductive mobilization of ribosomal labor away from other transcripts, and thus to futile fitness cost.

**Figure 6.**
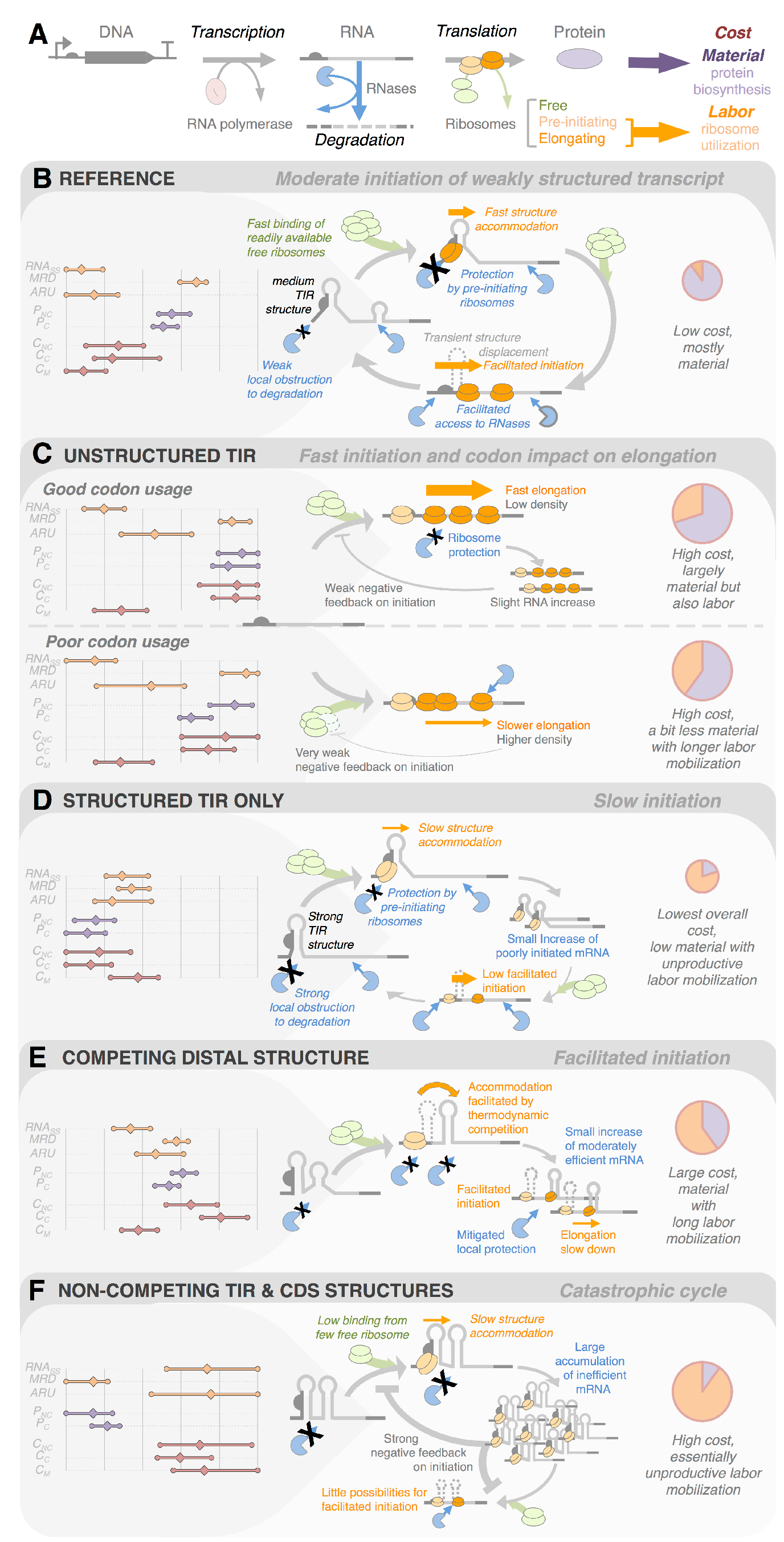
Impact of archetypal sequences on controlling translation efficiency and cost. **(A)** Key processes in the central dogma of molecular biology. This diagram introduces graphical elements used in the schematics below. **(BCDEF)** Combinations of sequence properties and consequent variations in translation efficiency determine diverse physiological costs. Left charts show the interquartile ranges of various measured phenotypic averages across factorial series, as shown (abbreviations as used throughout the text; the costs C_NC_, C_C_ and C_M_ are the opposite of W_NC_, W_C_ and W_M_, respectively). Schematic models in the middle provide a mechanistic framework for these data. Physiological consequences are rendered as schematic pie charts on the right, the size of which conveys the magnitude of the overall cost, while the slices differentiate material (violet) and labor (orange) contributions. Data are subset by combinations of categorical properties used for design as follow: **(B)** Medium STR_−30:+30_ ∩ Weak STR_+01:+60_ ∩ Weak STR_+31:+90_**(C)** *top:* Weak STR_−30+30_ ∩ WeakSTR_+01+60_ ∩High CAI; *bottom:*Weak STR_−30:+30_ ∩ Weak STR_+01:+60_ Weak CAI; **(D)** Strong STR_−30+30_ ∩ medium STR_+01:+60_ ∩ weak STR_+31:+90;_ **(E)** medium STR_−30+30_ ∩ medium STR_+01+60_ ∩ strong STR _+31:+90;_ **(F)** strong STR _-30:+30_ weak STR_+01:+60_ ∩ strong STR_+31:+90_.

## Discussion

Nearly half the ribosomes are dedicated to producing more ribosomes in growing *E. coli* cells (Li et al., 2014). This poses a compelling optimization problem, whereby ribosomes must be properly allocated to achieve the fine balance between ribosome production and that of other proteins necessary to maximize cell growth (Scott et al., 2014). Failure to do so can have lasting deleterious consequences on the cell fitness, which increases the value of ribosomes far beyond their production cost. Understanding how sequence properties influence this allocation pattern is essential from both fundamental and applied perspectives.

We precisely designed systematic sequence variations in the coding sequence of a particular reporter to dissect the phenotypic consequences while controlling, as much as possible, for statistical artefacts or confounding mechanisms. While we confirm that many known sequence features such as transcript secondary structure and codon usage bias are important, we also uncover hard-to-predict feedbacks between transcript degradation, translation and cellular homeostasis and demonstrate a large amount of remaining uncertainty in mechanisms that impact translational efficiency.

The three overlapping structures we explored (Figure 1C) may have two major functional consequences: i) intrinsic protection of the transcript from degradation; and/or ii) diminution of translation through decrease of initiation and/or elongation rates. Once recruited on transcripts, ribosomes can: i) sterically protect the stretch of mRNA they are sitting on; and ii) directionally unfold structures as they proceed along the transcript, thereby opening the way for immediately following ribosomes, as well as RNases. The interplay between these mechanisms gives rise to variety of archetypal phenotypic patterns that we detail below (Figure 6).

Sequences designed with mild TIR structure and no structures otherwise may serve as a baseline reference for discussion (Figure 6B). Medium TIR structures determine largely limiting initiation rates. These are nonetheless capable of driving sizeable protein production when free ribosomes are numerous enough to support another initiation before the TIR structure opened by the preceding event can refold, a process we refer to as ribosome cooperativity. The absence of strong downstream structures and the regular ribosome flow ensure basal rates of transcript decay. The low physiological charge associated with this configuration is mostly attributable to the material cost of moderate protein biosynthesis. This cost decreases when cells are grown in less productive conditions (see C_M_ and Figure 2M). The cost of ribosome utilization is minimal because initiation is fast enough to prevent transcript accumulation (low RNA_SS_), but slow enough to avoid ribosomal jams during subsequent elongation (medium MRD). The apparent ribosome utilization (ARU = MRD × RNA_SS_) is then kept at its lowest, ensuring efficient replenishment of the free ribosome pool and subsequent redistribution toward other transcripts. are limited by elongation rather than initiation. In the absence of other downstream structures, elongation rates are mostly dependent on codon usage (Figure 6C). Codon-adapted sequences support increased P_NC_ and even more P_C_. Although translation often favors transcript decay by disrupting otherwise stabilizing secondary structures, the situation is reversed under high initiation because translating ribosomes enter into kinetic competition with RNA_ses_ (Figure 4DE). While less codon-adapted transcripts exhibit higher MRD (Figure 5B), improved codon usage lead to higher RNAPTX and subsequent RNASS (Figure S5AB). Thus, punctual traffic jams at slowly translated sites lead to uneven ribosomal loads and provide more endonucleolytic opportunities than a smoother ribosome flow (Figure 6C). Ribosome protection has only a marginal impact on RNASS in our system and is most visible during the mRNA decay assay (Figure 4C and Figure 3E). This probably stems from our using a highly transcribed reporter, which likely limits the maximal initiation rate attainable through ribosome titration. We expect the importance of the above mechanisms to be amplified at lower transcription rates.

Under high initiation, codon adaptation is linked to a complex combination of signals, which altogether indicate smoother and more efficient elongation: increased protection, higher RNA abundance, lower ribosomal density and increased protein production. Our results supports the notion that strong codon biases in highly transcribed natural genes have been selected to minimize ribosomal retention on these transcripts (Kudla et al., 2009). Nevertheless, we only find a small positive association between CAI and growth (Figure 2J and S2IJ). Improving CAI should lead to increased fitness through lower ribosomal utilization—but would also incidentally augment ribosome protection, RNA abundance and protein production, ultimately resulting in higher material cost. The signal relating CAI to fitness might therefore be blurred by nearly compensatory transactions between these two cost components (Figure 6C).

Consider now transcripts with strong TIR, but no downstream, structures (Figure 6D). Pre-initiating ribosomes that bind these molecules at unstructured upstream sites must accommodate the folded TIR to eventually initiate translation. Structure accommodation is an active process that takes place at the ribosomal platform and may occasionally require specialized chaperones, such as the ribosomal protein S1 (Duval et al., 2013). Other regulators (*e.g.* S15) can instead stabilize these complexes, thereby trapping ribosomes in a lasting stalled state (Marzi et al., 2007). In either case, pre-initiating ribosomes remain bound to transcript for long periods of time, during which they are effectively sequestered away from other transcripts. Here, the bound upstream sequence and the structure being unfolded are likely protected, whereas the unstructured downstream sequence is fully accessible to RNAses. This determines moderate transcript accumulation and medium ARU. Subsequent decrease in free ribosomes contributes to lower effective initiation rate through hindering timely ribosome cooperation. Given the low resulting protein production, this initiation-based ribosome sequestration must determine most of the small fitness cost measured under regular growth condition. This cost is markedly aggravated under increased ribosome limitation (C_M_ and Figure 2M).

The accommodation of TIR structures may be thermodynamically facilitated when associated with stronger structures overlapping downstream (Figure 6E). Subsequent improvements of initiation are, however, likely mitigated by lower elongation rates through that distal structure. Although moderately translated individually, such transcripts comprise enough structures to cumulate to intermediate level. They collectively yield sizeable protein production, though at a relatively high cost, due to high ARU. In isolation, protein production data suggested that strong distal structures could determine increased translation efficiency (companion paper). Integrative analysis instead reveals individual inefficiency and a collective performance driven by a moderate though deleterious transcript accumulation (Figure 2C and Figure 3F).

In the worst case, distal structures are stably associated with non-overlapping TIR structures (Figure 6F). This configuration describes poorly initiated and intrinsically stable transcripts. The consequent rarity of translation minimizes structure unfolding within the coding sequence (lowest MRD) and thus ensures maximal transcript stabilization (highest RNA_SS_). Titration of ribosomes by cumulating transcripts further limits the opportunities for cooperative initiation at the structured TIR (largest ARU). These interdependencies determine a catastrophic cycle of ever increasing unproductive ribosome utilization, global initiation decrease and augmented stabilization that eventually leads to extreme and futile labor costs. The absence of protein production from many such pathologically abundant transcripts highlights the tightness of the ensuing initiation lock. Subtle parameter variations departing from this situation can account for the continuum of phenotypes in our experiments. For example, slight improvement of initiation rates may lead to moderate protein production at large fitness cost if the transcripts remain stable enough to accumulate; or to lower fitness cost with little increase in protein production if, in contrast, transcripts are heavily destabilized.

The complex functional interplay of competitive structures along the transcript can be exploited for regulation, as in the mechanisms underlying riboswitches. In these cases, it is likely optimal to minimize ribosome sequestration by maintaining low transcript abundance and using an efficient RBS. Such a configuration, however, is prone to expression noise which itself may be non-optimal. When there is little TIR structure, keeping RNA high and initiation medium has recently been found to be relatively optimal (Ceroni et al., 2015). The advantage of medium initiation might arise from then lower ribosome sequestration on *de facto* poorly elongated heterologous transcripts despite the intended codon-optimization. Our model suggests that improving the initiation-elongation balance should improve both protein production and translation capacity in that case.

Our analyses suggest that successful low-cost heterologous gene expression should rely on well-initiated transcripts exempt of secondary structures in a TIR region spanning ~50 nts on either side of the start codon. Engineering a medium-strength structure immediately downstream of that window should impact initiation favorably by competing with unexpected TIR structure. Over-strong structure there and elsewhere in the downstream sequence can lead to transcript stabilization, slower elongation and unnecessary fitness costs. Codon usage should be optimized, but not take precedence over structure optimization. Transcription rates should be optimized, most especially in structured sequences, since high RNA abundance can uselessly sequester ribosome, potentiate the effect of secondary structures, modify the global translation regime of the cell and eventually impact fitness without necessarily improving production.

The production of ribosomes is tightly regulated. Antibiotic-mediated inhibition of translation promotes rRNA transcription in *E. coli*, possibly through mitigation of the stringent response driven by decreased protein biosynthesis and subsequent amino-acids accumulation (Scott et al., 2010). We expect a similar response to genetically induced perturbations that lead to sequestering of existing ribosomes. Increasing rRNAs are expected to up-regulate ribosome biogenesis (Paul et al., 2004), with potentially ambivalent consequences: although that could eventually improve the cell‘s translation capacity, it would momentarily mobilize dwindling ribosomes at the risk of strengthening possible deleterious feedbacks. Additional small-scale experiments on exemplary strains will be necessary to determine in which circumstances these responses are beneficial.

Normalization of protein by mRNA abundance is often used as a proxy for translation efficiency—strictly defined as the current of ribosomes over a transcript. This ratiometric approach controls for the dependence of mRNA decay and abundance on translation rate (Li, 2015). Our results suggest that the converse relationship must also be considered. Translation efficiency is non-linearly dependent on transcript abundance, because: i) a finite number of ribosomes must be allocated to a large excess of mRNAs; and ii) their relative availability modulates the cooperative initiation of TIR-structured transcripts. A measured protein/mRNA ratio is therefore contingent on particular conditions and not necessarily operational in other contexts. For instance, increasing the transcription of a poorly initiated construct can instigate the aforementioned catastrophe and a dramatic drop of the protein/mRNA ratio.

The above makes clear that the complex and pleiotropic roles of RNA structures and their implications in intricate feedbacks among initiation, elongation and degradation are key to understanding physiologically efficient protein production. However, current predictions of these features and their dynamics are not accurate enough to support fully operational models (see companion paper). Other unknown sequence properties and non-linear behaviors are likely to limit our predictive power as well. As a suggestive illustration, the 56 replicate factorial series studied in this work were designed to systematically explore the exact same properties space but often show widely divergent average phenotypic responses from one another (Figure 7A). For examples: series #26 and #205 determine high RNA_SS_ and subsequent low growth, but only the former show defect in protein production; series #56 and #71 show weak protein production improvement under coupling, whereas #49 instead enjoys a particularly marked increase; #227 and #2 produce low protein in both conditions, but while the former shows high growth in all conditions, the latter does not but nonetheless experiences large rescue under coupling (Figure 7B). The list of such idiosyncrasies is vast. Further work is needed to capture the details enabling reliable prediction from sequence alone. We have hope that this dataset will enable others to advance in that direction and inspire other endeavor in this area.

**Figure 7.**
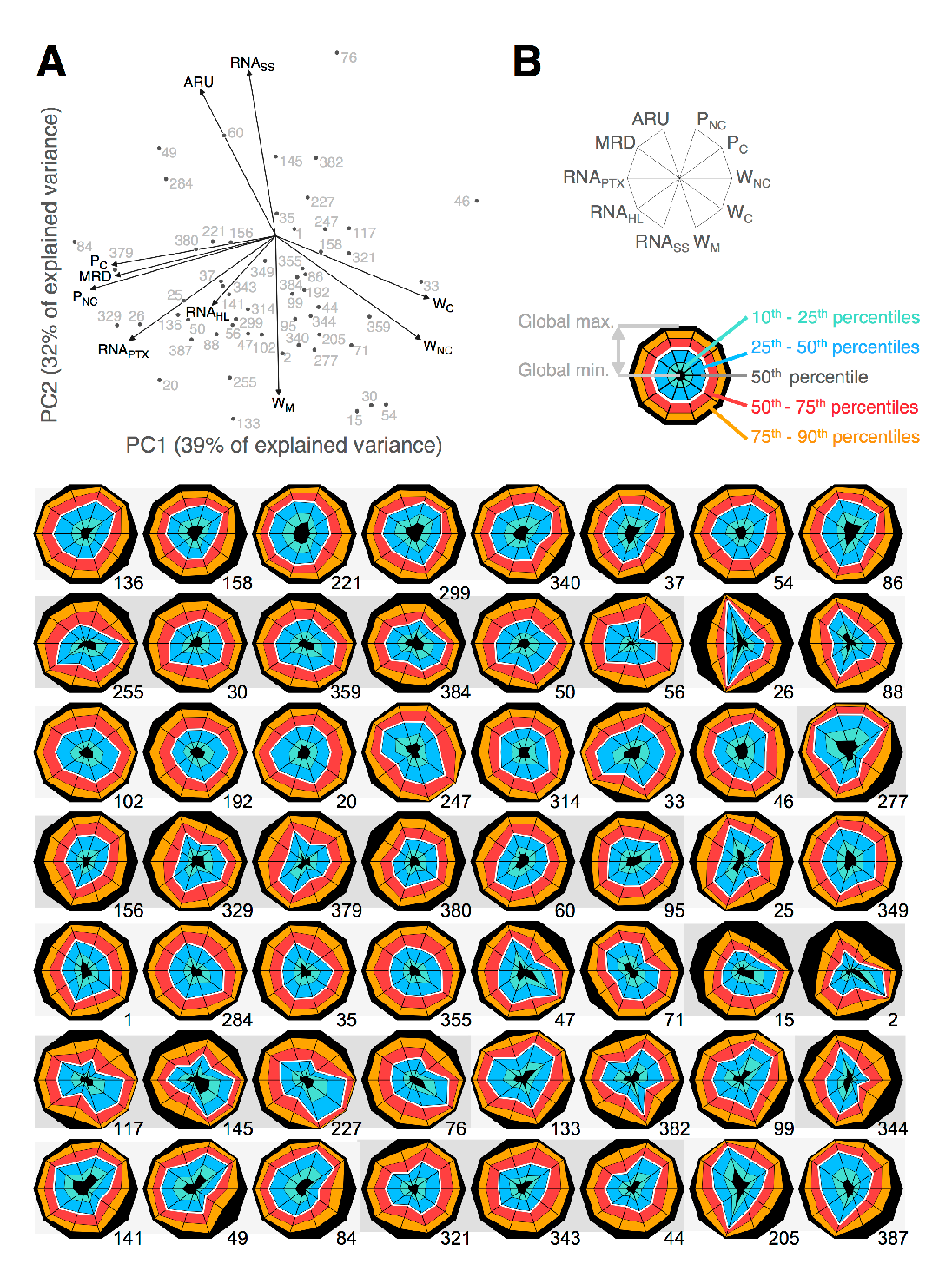
Extensive phenotypic diversity between replicate factorial series. **(A)** Principle component analysis highlights the phenotypic spread of factorial series. The analysis is based on the correlation matrix between the series-wise means of shown phenotypic variables. **(B)** Visualization of the ensemble phenotypic differences between series using spider plots. Although each series explores the same property space, small initial phenotypic differences may cascade into the observed diversity. Understanding these differences represents a challenge in predictive biology.

## AUTHOR CONTRIBUTIONS

GC conceived the work; GC performed the experiment and processed the data; GC and APA analyzed the data; GC and APA wrote the manuscript.

## ACKNOWLEDGMENTS

We thank Premal Shah, Joshua Plotkin and Luca Ciandrini for useful discussions. GC was funded by the Human Frontier Science Program (LT000873/2011-L). We acknowledge financial support by the Synthetic Biology Engineering Research Center (SynBERC). This work used the Vincent J. Coates Genomics Sequencing Laboratory at UC Berkeley (NIH S10 Instrumentation Grants S10RR029668 and S10RR027303).

